# Human wastewater contamination drives the emergence of novel multidrug resistant bacteria in the Galápagos marine ecosystem

**DOI:** 10.1101/2025.09.12.675863

**Authors:** Arnav Lal, Jade C. Riopelle, Katherine Villarin, Maya Mathur, Lia Enriquez, Rui Xiao, Naomi Phemister-Jimenez, Katherine Gilbert, Stephen D. Cole, Maddie Tilyou, Kelly P. Kennedy, Ernesto Vaca, Wilson Castillo, Michael Weisberg, Lisa M. Mattei, Daniel P. Beiting

**Affiliations:** Department of Pathobiology, School of Veterinary Medicine, University of Pennsylvania, Philadelphia, Pennsylvania, 19104, USA; Harvard Medical School, Boston, Massachusetts, 02115, USA; Galápagos Education and Research Alliance, Penn Global Research Institute, University of Pennsylvania, Philadelphia, Pennsylvania, 19104, USA; Department of Philosophy, School of Arts and Sciences, University of Pennsylvania, Philadelphia, Pennsylvania, 19104, USA; Perry World House, University of Pennsylvania, Philadelphia, PA, 19104, USA; Parque Nacional Galápagos, Isla San Cristóbal, Galápagos, Ecuador

**Keywords:** Galápagos, metagenomics, wastewater, antimicrobial resistance, multi-drug resistance, nanopore sequencing, surveillance, One-health, community science

## Abstract

Since the publication of Charles Darwin’s *On the Origin of Species* in 1859, the Galápagos archipelago has become emblematic of natural selection and evolution. While the lens of evolution in the Galápagos has traditionally focused on iconic megafauna, including finches, marine iguanas, and giant tortoises, the marine environment is also home to diverse microbial ecosystems that are constantly evolving under selective pressure from environmental factors such as human activity. We focused on the second most populated island within the archipelago — San Cristóbal — an island that has experienced rapid urbanization and intense international tourism pressure. Using a ‘lab-free’ approach, we spatiotemporally mapped wastewater contamination around San Cristóbal. On-site metagenomic sequencing revealed a stark shift in genera and a higher count of antimicrobial resistance genes at wastewater-associated sites. Over 40% of lactose-fermenting *Enterobacteriaceae* isolates collected from sewage and wastewater outfall exhibited multidrug resistance (MDR). Long-read sequencing and *de novo* assembly of bacterial genomes and plasmids from MDR *Escherichia coli* revealed frequent and rapid reassortment of antimicrobial resistance genes on plasmids, generating unique antimicrobial resistance profiles. This study not only provides a framework for conducting antimicrobial resistance research in low-resource settings but also underscores the impact of wastewater contamination on the environmental AMR landscape and highlights potential threats to human and animal health.

## Introduction

Antimicrobial resistance (AMR) is a major threat to global public health. Excluding *Mycobacterium* tuberculosis, bacterial AMR resulted in roughly 5 million deaths globally in 2019^1^ – accounting for 9% of all deaths worldwide – and this number is forecasted to double by 2050^2^. Lower- and middle-income countries (LMICs) are disproportionately affected by AMR^1^ due to myriad factors, including inadequate surveillance and waste management infrastructure, lower vaccination rates, rapidly developing agricultural practices, increased urbanization, and antimicrobial misuse^3–6^. Intensified use of antimicrobials during the COVID-19 pandemic^7^, together with the continued widespread use of antimicrobials in livestock globally^8^, increases the likelihood of higher morbidity and mortality from AMR. Despite the global prevalence and importance of AMR, the extent and mechanisms of AMR dissemination in the environment are poorly understood. This knowledge gap is particularly severe in LMICs, where a lack of research laboratory infrastructure hinders surveillance efforts, AMR phenotyping, and genomic identification of AMR genes.

The Galápagos archipelago is an ideal natural laboratory in which to study the impact of expanding human activity on the emergence of antimicrobial resistance in marine ecosystems. Although the majority of the archipelago’s islands remain uninhabited, four have a small but growing human population, with the majority settled on the islands of Santa Cruz and San Cristóbal (approximately 17,000 and 8,300, respectively, as of the 2022 census^9^). Both islands have experienced rapid urbanization with annual population growth rates of 6%, three times the rate of mainland Ecuador, and both are the focus of intense pressure from land-based international tourism. In 2022, for example, San Cristóbal saw more than 76,000 tourist visits to areas concentrated within a 2.5 km area around Puerto Baquerizo Moreno, the island’s largest town^10,11^. International tourism likely increases both the diversity of strains introduced to the environment and the variety of antibiotic exposure. Furthermore, inadequate wastewater management on San Cristóbal has been highlighted as a source of fecal coliform contamination and AMR in recreational water^12,13^. Altogether, the rapidly growing population and dynamic flow of tourists, alongside the high density of marine mammals that frequent these islands, raise the potential for AMR interchange between diverse human-human and human-animal hosts within the marine environment, thus imperiling One-Health in one of our planet’s most cherished ecosystems. Despite these unique circumstances, little is known about the impact of wastewater contamination on the Galápagos’ environment. A more comprehensive view of how human activity impacts the development of AMR in this marine ecosystem requires spatial and temporal mapping of wastewater contamination in parallel with untargeted metagenomics and genome sequencing to identify relationships between bacterial taxa, AMR genes, and resistance phenotypes.

To address this knowledge gap, we developed a mobile microbiology laboratory to identify ‘hot spots’ of human wastewater contamination in marine environments around the island. Metagenomic sequencing of marine water at these hot spots identified human fecal bacteria and AMR gene profiles that matched what was observed contemporaneously in local municipal sewer systems. Moreover, lactose-fermenting *Enterobacteriaceae* selectively cultured from contaminated marine environments frequently exhibited multidrug resistance (MDR) phenotypes, and whole genome sequencing of *Escherichia coli* strains revealed rapid and frequent reassortment of AMR genes within plasmids, ultimately giving rise to potentially novel MDR bacteria. Taken together, these data underscore the need for environmental AMR surveillance and risk assessment in the Galápagos archipelago to protect local human populations, tourists, and the marine ecosystem.

## Results

### Geospatial mapping of human fecal contamination in the marine ecosystem around San Cristóbal, Galápagos

Traditional microbiology and microbial genomic laboratories occupy a large footprint and are highly dependent on a reliable cold chain and continuous electrical supply. Most research instrumentation is not meant to operate outside of this environment, and consumables such as media plates and plasticware may not be readily available in lower- and middle-income countries (LMICs). These factors strongly restrict locations where robust environmental surveillance of AMR can occur. To circumvent these issues, we developed and optimized a portable microbiology field laboratory **(Supplementary Table 1)** that allowed us to detect human fecal contamination in marine ecosystems **(Figure 1A)**, isolate bacterial strains for AMR profiling **(Figure 1B)**, and carry out untargeted metagenomic sequencing and whole genome sequencing of bacterial isolates **(Figure 1C)**.

**Figure 1:**
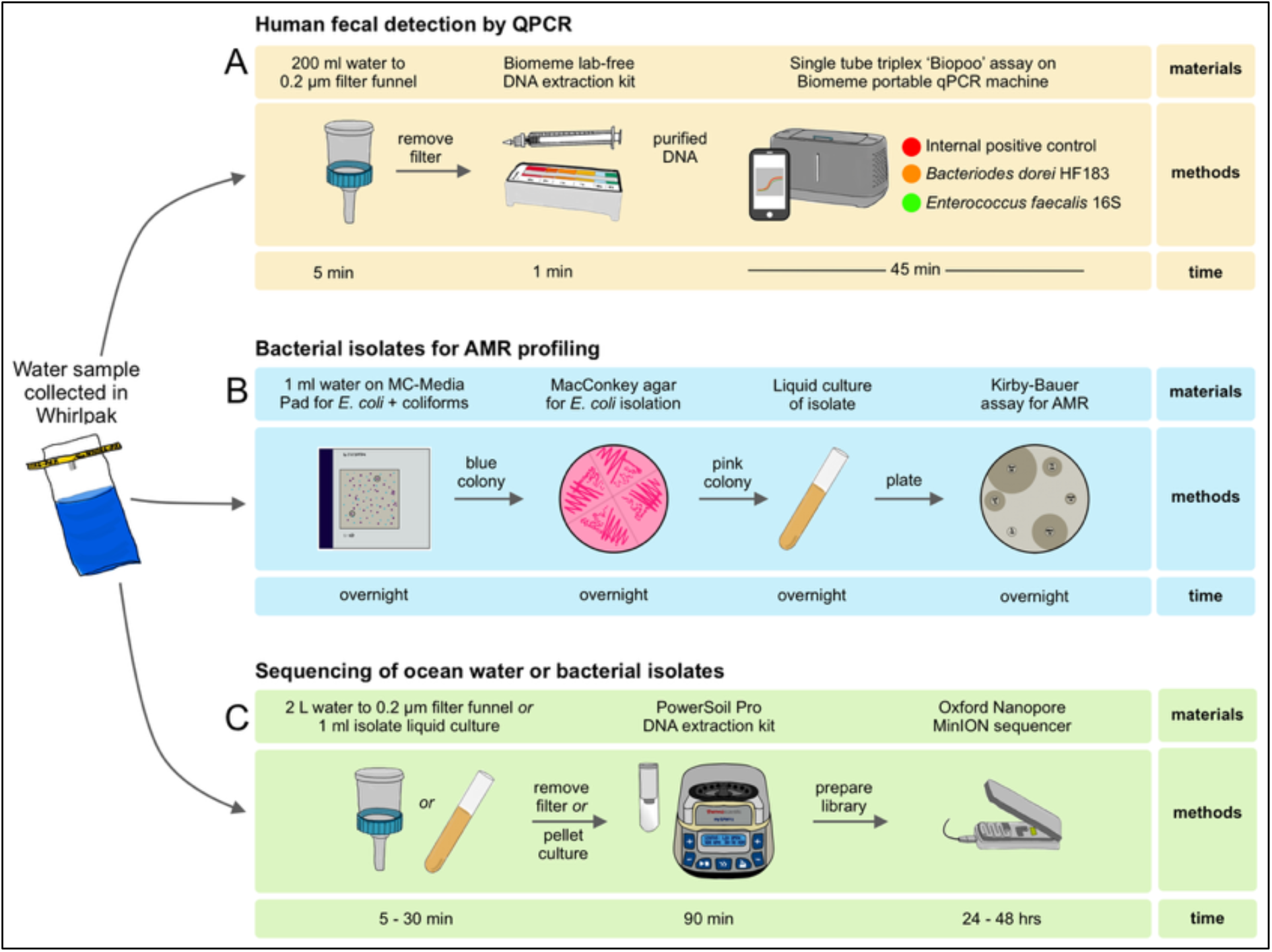
A ‘lab-free’ and portable AMR discovery platform. Ocean water samples collected in sterile bags were processed through one or more of three workflows, each of which emphasized field-ready assays and portable equipment: (A) QPCR for the detection of human fecal contamination. (B) AMR profiling of *E. coli* isolates from marine ecosystems or municipal sewer systems. (C) Long-read sequencing of ocean water, wastewater, or *E. coli* isolates.

To identify enteric pathogens in the marine ecosystem surrounding San Cristóbal, samples were collected from 16 marine sites, two freshwater sites, and two municipal sewage sites (**Supplementary Table 2**) over a two-year period. These samples were evaluated using a field-ready lyophilized triplex qPCR assay that distinguishes human fecal contamination from other fecal sources using the HF183 marker gene derived from *Bacteroides dorei* 16S rRNA **(Figure 1A)**^14^. Sample sites were dispersed around the island **(Figure 2A)** with the highest density of sampling occurring within 1.5 km of the center of Puerto Baquerizo Moreno **(Figure 2B)**, proximal to the human population and tourism epicenter. Four sites (Tongo Reef, Playa Lobería, Playa Baquerizo, and Playa Mann) were regularly frequented by locals and tourists and were located within 3 km of the town center. In contrast, four other sites (León Dormido, Puerto Chino, Bahía Rosa Blanca, and Punta Pitt) were distant tourist locations situated more than 15 km from the town center, and one site (Isla Española) was located off the coast of an unpopulated island approximately 50 km south of San Cristóbal (**Figure 2A**).

**Figure 2:**
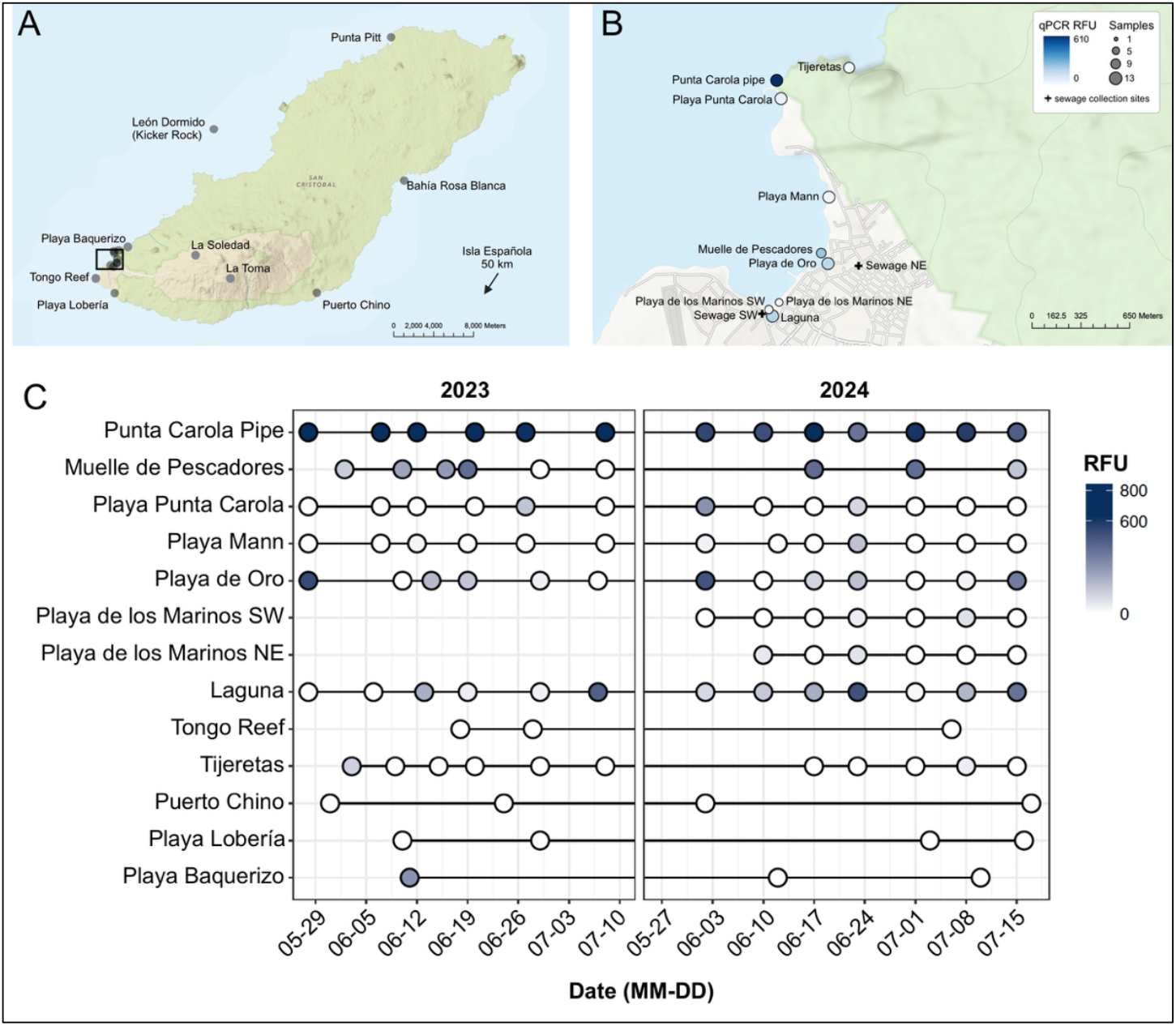
Geospatial mapping of human fecal contamination in the Galápagos marine ecosystem. (A) Map of San Cristóbal island. Points indicate locations where water was collected and assayed for fecal contamination by qPCR. Box indicates the region shown in (B), where sampling was most intense and which is centered on the Puerto Baquerizo Moreno where population density and tourism are highest on the island. (C). Dot plot showing relative fluorescent units (RFU) results for qPCR assay over a two-year period for 13 sites shown. Each point represents a single sampling event. Maps were created in ArcGIS. Detailed information and location data for sampling sites can be found in **Supplementary Table 2**.

All samples tested positive for the pan-stool marker by qPCR. Positive pan-stool markers and negative human-stool markers suggest the presence of animal fecal contamination, consistent with the observation that all sites sampled are frequented by marine species, most notably sea lions, marine iguanas, and a diverse array of fish and avian species. Multiple sites also tested positive for the human-specific stool marker, likely due to wastewater contamination. We observed spatial **(Figure 2B)** and temporal **(Figure 2C)** variability in human fecal contamination, with the strongest and most consistent signal observed at the Punta Carola pipe, a site that has been locally confirmed and previously reported to be a point of sewage outfall^12^. Other sampled sites demonstrated lower and/or more variable signals for human fecal contamination by qPCR. For example, at the Muelle de Pescadores site, a fishing pier with frequent boat docking where residents occasionally swim, human fecal contamination was observed during both summers of data collection, apart from the month of July 2023. At Playa de Oro, a sea lion-inhabited beach with reported wastewater outfall, high and low human fecal qPCR findings can be seen throughout both summers. This same pattern was observed at the Laguna, a brackish water site where wastewater outfall has also been reported. While other sites demonstrated occasional detection of human fecal contamination, they rarely reached the levels of the Carola pipe, Playa de Oro, the Muelle de Pescadores, or the Laguna. Notably, proximity to a site with a strong qPCR signal for human fecal contamination did not guarantee a positive assay. For example, the strong and consistent positive tests at Punta Carola pipe were not reflected at Playa Punta Carola, perhaps due to the presence of an extended rocky breakwater between the sewage outfall and the beach area **(Figure 2B)**. Additionally, the two sampling sites at Playa de los Marinos are separated from the Laguna by a small beach strip that likely prevents coliform movement during times of low precipitation.

The positive qPCR assays for human stool markers at multiple sites raise concerns about live human fecal contamination in waters, especially near the port town. Notably, the sites with the highest signals by qPCR are all locations with suspected (Playa de Oro, Muelle de Pescadores, and Laguna) or documented (Punta Carola pipe) wastewater outfall. Our qPCR assay detects DNA and thus cannot distinguish between live or dead fecal bacteria. To determine whether the locations identified by our qPCR testing contain live fecal bacteria, we evaluated *E. coli* and coliform growth directly from ocean water using MC-Media pads, a simplified, miniaturized, and portable alternative to traditional petri dishes that provide robust coliform and *E. coli* CFU counts **(Figure 1B)**. This analysis revealed >3000 CFU/mL of coliforms from the water around the Punta Carola pipe and occasionally similarly high values at the Laguna site, indicating that our qPCR assay results reflect contamination with live fecal coliforms **(Supplementary Figure 1)**.

### Metagenomic sequencing of marine water identifies common human fecal contaminants linked to wastewater infrastructure

Although our data highlight potential sites with wastewater contamination, these approaches do not reveal the broader impact of wastewater on marine microbial diversity, nor enable a detailed view of microbial gene presence, such as AMR genes. Thus, we conducted long-read metagenomic analysis of water collected from selected sites across the town and the broader archipelago **(Figure 1C)**. Since wastewater has been shown to reflect the enteric microbiome of resident human populations^15^, we also collected samples of untreated sewage from two sites within the town. Over the summers of 2022 and 2023, we sampled eight sites, two of which were sampled in both years. Sites clustered predominantly into two groups based on taxonomic abundance of genera (**Figure 3A, top dendrogram**). One group includes a remote beach (Playa Baquerizo), a distant uninhabited island (Española), a commonly used beach within city limits (Playa Mann), and the port’s main fishing pier (Muelle de Pescadores) in 2023. Apart from the pier and Playa Mann, all sites are relatively remote, and the genera observed at these sites predominantly include *Synechoccous, Sphingobacterium, Prochlorococcus*^16^, *Chaetoceros*^17^, *Ostreococcus*^18^, *Planktomarina*^19^, *Pseudoalteromonas*^20^, *Alteromonas*^21^, and *Vibrio* strains, which are associated with marine and environmental sources^22–24^. The relative paucity of enteric microbes detected at these sites suggests little or no exposure to wastewater outfall. Interestingly, the fact that one of the most central beaches (Playa Mann) – a site heavily visited by both locals and tourists – has a taxonomic profile similar to remote sites **(Figure 3A)** and shows limited detection of human fecal contamination by qPCR **(Figures 2B and 2C)** suggests that recreational activity in and around the water is not sufficient as a source of fecal contamination.

**Figure 3:**
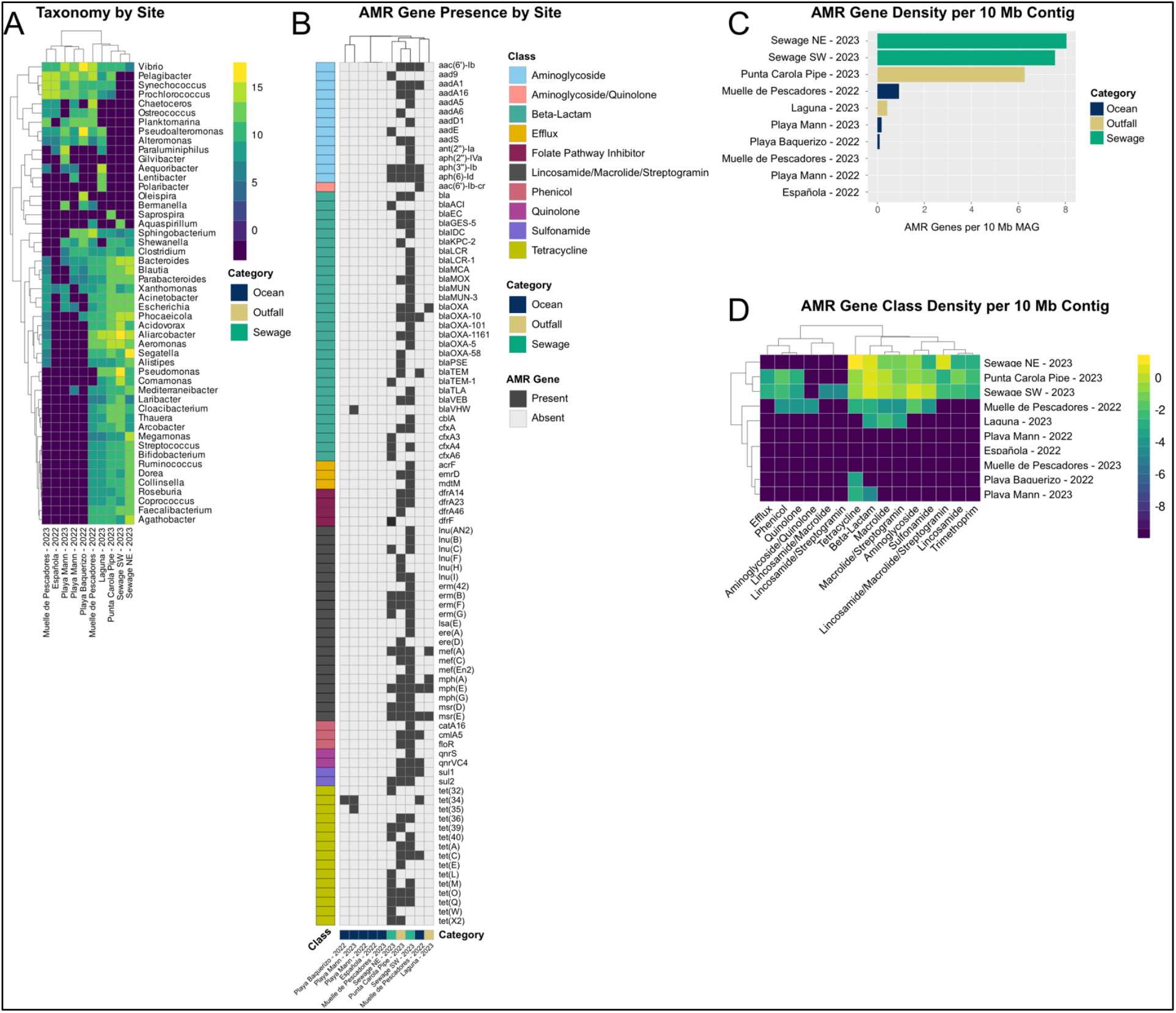
Metagenomic sequencing identifies shifts in the marine microbiome at sites of fecal contamination. (A) Heatmap showing relative taxonomic abundance (Log2 bases per million) at the genus level for bacteria and archaea in water and sewage metagenome samples, as classified by the CZ ID tool^25^. Only taxa with >85% nucleotide identity and >500 bp alignment length are included. The top 50 most abundant taxa are shown. (B) Presence/absence results for specific antimicrobial resistance (AMR) genes detected in contigs >500 bp using AMRFinderPlus^26^. Columns are clustered by average linkage hierarchical clustering using Jaccard distances. Colors to the left and below indicate AMR gene class and sampling site, respectively. The total AMR gene density (C), and the AMR density broken down by AMR gene resistance class (D) are displayed as the number of AMR genes per 10 Mb of assembled contig length.

The second cluster (**Figure 3A, top dendrogram**) is comprised primarily of wastewater-associated sites, including the two sewage samples, the two sites where wastewater outfall is known to occur (Punta Carola pipe and Laguna), and the fishing pier (Muelle de Pescadores) in 2022. A very different core set of genera predominates at these locations, including *Streptococcus, Bifidobacterium, Arcobacter, Pseudomonas, Clostridium, Bacteroides, Acinetobacter*, and *Escherichia*. Many of the genera detected in these samples are potential pathogens of both humans and animals, and the presence of *Escherichia* indicates coliforms within the outfall and, not surprisingly, the municipal sewage system. Interestingly, the impact of wastewater contamination can be dynamic, as observed with Muelle de Pescadores, where the metagenomic profile changed substantially, moving from the cluster of wastewater-associated sites in 2022 to the cluster of more remote sites in 2023. The similarity in taxonomic profiles between wastewater outfall sites and municipal sewage sites, together with our qPCR **(Figure 2)** and CFU results, strongly supports the conclusion that these locations are contaminated with wastewater.

### AMR genes are detected at wastewater-associated sites

Since human waste can be a source of AMR in the environment^27–29^, we next sought to use our long-read metagenomic sequencing data to identify the presence of AMR genes at each site as a proxy for environmental AMR burden. We observed a large and diverse set of AMR genes across our metagenomic samples, including 94 resistance genes spanning nine antimicrobial classes (**Figure 3B**). The highest density of AMR genes was observed in contigs generated from sewage samples, followed by the Punta Carola outfall site, while other sites contained few or no AMR genes (**Figure 3C**). Separation of the AMR gene density by resistance class highlights a clear pattern of similarity between the Punta Carola pipe and the two sewage sites (**Figure 3D**), including a relatively strong presence of tetracycline, beta-lactam, and macrolide AMR genes, among others. A lower but detectable level of AMR genes in similar classes was observed in the 2022 Muelle de Pescadores sample (but notably not the 2023 sample), along with the sampling of the Laguna. Taken together, these data link the wastewater-driven taxonomic shift in the marine ecosystem to a high environmental burden of AMR potential.

### Assessment of antimicrobial-resistant *E. coli* from marine and wastewater environments

The presence of AMR genes does not necessarily reflect the actual ability to resist antimicrobials^30,31^. Therefore, we next set out to determine the AMR phenotype of bacteria recovered from the marine ecosystem around the island. For this analysis, we chose to focus on lactose-fermenting *Enterobacteriaceae* since *Escherichia* was detected by metagenomic sequencing at all five wastewater-associated sites (**Figure 3A**), and because of the relative ease with which they can be isolated by culture and their potential to act as pathogens in humans and animals. In total, 183 lactose-fermenting *Enterobacteriaceae* isolates, predominantly *E. coli*, were cultured directly from ocean water (see methods) and subsequently assessed for antimicrobial resistance against a panel of clinically relevant antibiotics through Kirby-Bauer disk diffusion assays **(Supplementary Table 3)**. For every antibiotic examined, the proportion of isolates showing a resistant phenotype was similar between municipal sewage and marine sites with suspected or confirmed wastewater outfall (**Figure 4A**). Streptomycin represented the antibiotic class with the highest overall resistance (86 of 183; 47%), followed by Ampicillin (74 of 183 isolates; 40.4%).

**Figure 4:**
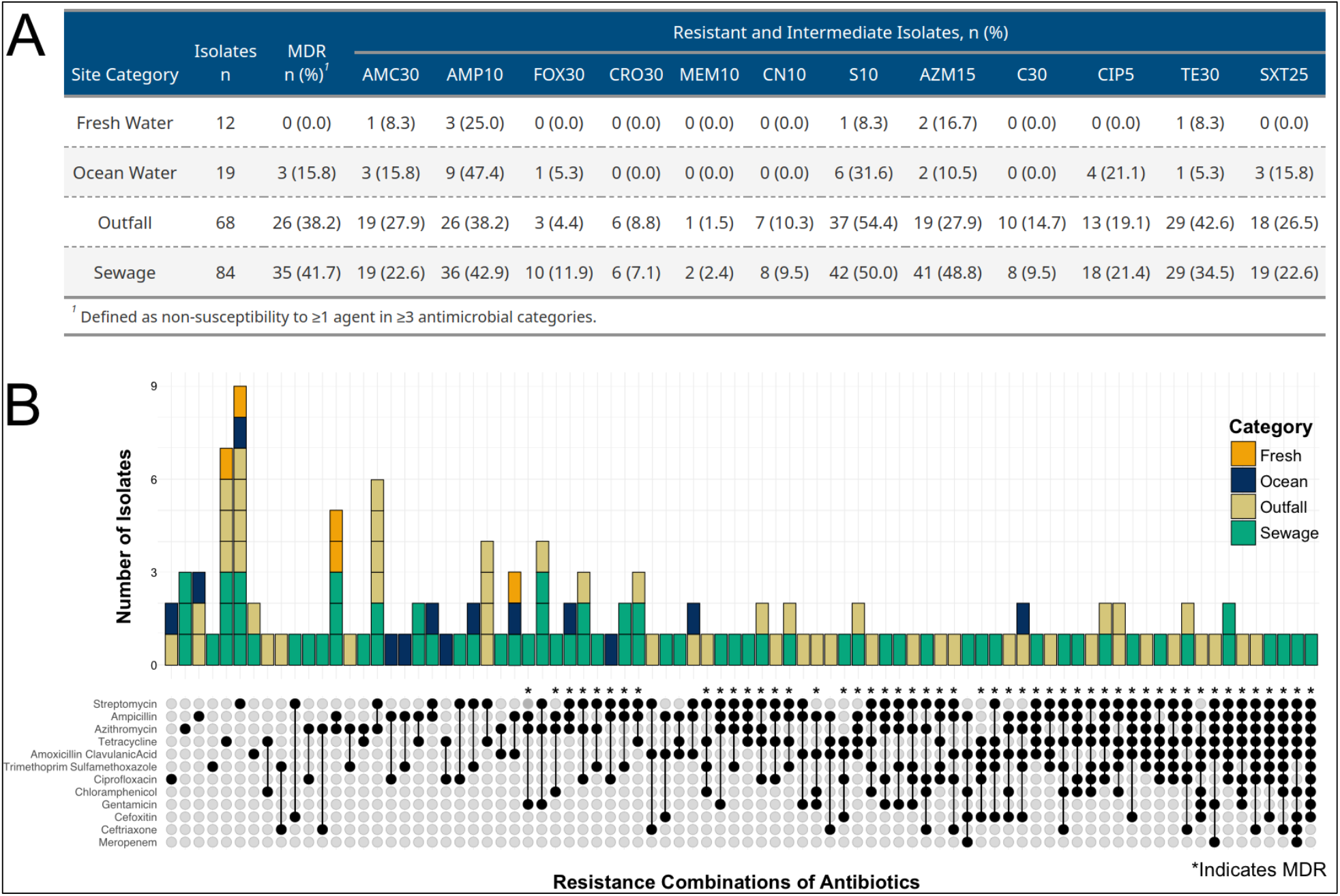
Antimicrobial resistance profiles of *E. coli* isolates. (A) Table showing counts of *E. coli* isolates exhibiting non-susceptibility to antibiotics, as assessed through standardized Kirby-Bauer disk diffusion assays, separated by site classification of fresh water, ocean water, outfall, or sewage. Multidrug-resistant (MDR) counts are defined as resistance to three or more different antibiotic classes. Antibiotic classes are abbreviated as follows: AMC: amoxycillin/clavulanic acid, AMP: ampicillin, AZM: azithromycin, C: chloramphenicol, CIP: ciprofloxacin, CN: gentamicin, CRO: ceftriaxone, FOX: cefoxitin, MEM: meropenem, S: streptomycin, SXT: trimethoprim-sulfamethoxazole, TE: tetracycline. (B) Stacked bar chart (top) and ‘Upset’ plot (bottom) showing number of isolates and category for different antibiotic resistance combinations. Asterisks indicate AMR combinations that meet the criteria for being multidrug resistant (MDR).

Multidrug resistance (MDR), defined as resistance to antibiotics from three or more different classes, was only observed in three isolates recovered from non-wastewater sites. In contrast, 26 of 68 isolates (38.2%) from wastewater outfall samples and 35 of 84 (41.7%) sewage samples were MDR. Classifying all 183 isolates by their resistance profiles highlights 64 MDR isolates that gave rise to 50 unique resistance profiles (**Figure 4B, asterisks**). While ampicillin and tetracycline resistance were present in most MDR isolates, the large number of unique MDR combinations hints at a potential genetic diversity that cannot be explained solely by the transmission of one or a few independent resistant *E. coli* lineages. Some sewage and outfall isolates are phenotypically resistant to six or more classes of antibiotics. Taken together with our metagenomic data **(Figure 3)**, these results highlight a diverse and functional AMR landscape around the island. Notably, given the classification of carbapenem-resistant *Enterobacterales* (including *E. coli*) as the most critical priority by the WHO^32^, we observed three unique strains carrying phenotypic resistance to meropenem even within this remote island.

### Phylogenomic analysis of antimicrobial-resistant *E. coli* highlights plasmid-localized AMR genes as a potential driver of multidrug resistance

The presence of AMR – particularly MDR – in sewage, outfall, and to a lesser extent in ocean water around San Cristóbal prompted us to explore the genomic basis of these resistance phenotypes within *E. coli*. We selected 53 of the 183 lactose-fermenting *Enterobacteriaceae* isolates with AMR phenotypes **(Figure 4)** for long-read sequencing, focusing primarily on MDR isolates **(Supplementary Table 3)**. *De novo* assembled contigs were evaluated based on size and GC content, resulting in the removal of eight samples with potential poor quality (e.g., co-assemblies of multiple or non-*E. coli* genomes, incomplete sequences, etc.) **(Supplementary Figure 2 and Supplementary Table 4)**. Of the remaining 45 high-quality assemblies, all yielded complete, ungapped bacterial genomes, and 37 included at least one complete plasmid **(Supplementary Table 4)**. Using these genomes and plasmids, we broadly classified our *E. coli* into clades based on Clermont typing^33^ and multi-locus sequence typing (MLST)^34^. This analysis identified five distinct phylogroups in our study, including: 21 group A isolates (commensals associated human and animal colon); 15 group B1 isolates (environment- and animal-associated); one group B2 isolate (highly virulent, human-associated); five group D isolates (more virulent, extraintestinal pathogen); two group E isolates (related to D and associated with foodborne outbreaks but can be commensal); and one group G isolate (similar to B2, human- and animal-associated, potential zoonotic transmission) **(Figure 5)**. Notably, our analysis identified several globally distributed, high-risk pandemic clones associated with MDR, including ST10, ST58, ST69, and ST155 (**Figure 5**)^35^.

**Figure 5:**
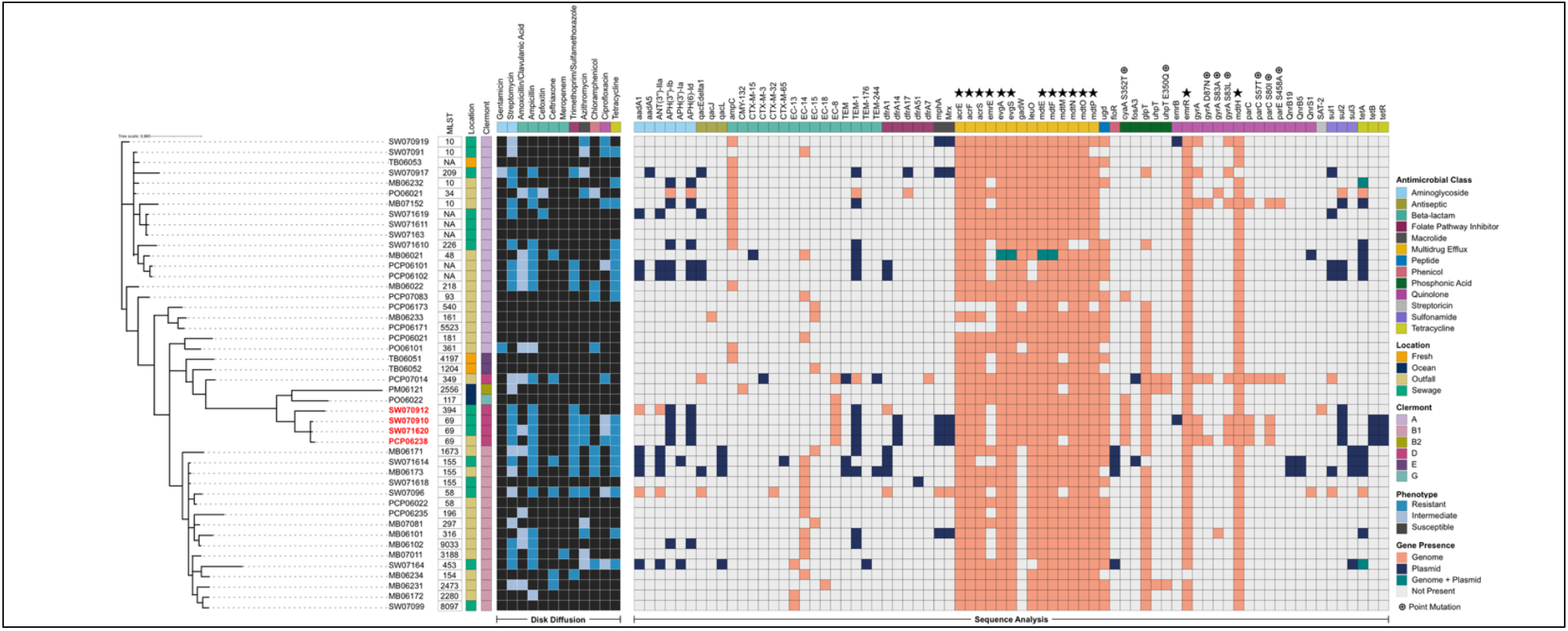
Integrating AMR phenotypes and genotypes of *E. coli*. Visualization showing AMR genes (top) detected in either the genome (pink) or plasmid (dark blue) or both (green) for 45 E. coli isolates (left). White cells indicate no AMR gene detected. Isolates were clustered based on single-copy genes (phylogenetic tree) and grouped by class of AMR (top; color). Kirby-Bauer disk diffusion assay results are shown (dark heatmap, left). Stars indicate AMR genes observed mainly in the genome (top). Circled cross symbol (top) indicates AMR genes with point mutations. Red labeled isolate identifiers (left) indicate multidrug resistant Clermont D group *E. coli* described in more detail in **Figure 6**.

To complement our phylogenetic analysis and link AMR phenotypes to specific genes, we annotated AMR genes on all isolated chromosomes and plasmids (see methods) (**Figure 5**). These analyses identified multiple genes and operons, including multidrug efflux genes and quinolone resistance genes such as *acr(E/F/S), mdt(E/F/H/M/N/O/P), emr(E/R)*, and *evg(A/S)*, that were present on the chromosome of nearly every isolate examined and rarely, if ever, detected on plasmids **(Figure 5, top, stars)**. Aside from these examples, most AMR genes we identified were either never or only rarely found on the chromosome but instead were localized to a bacterial plasmid. For example, AMR genes belonging to aminoglycoside, tetracycline, sulfonamide, or macrolide classes were almost exclusively plasmid localized **(Figure 5, top)**, thus highlighting the importance of long-read sequencing and *de novo* plasmid assembly for AMR gene discovery. Our analysis also identified a subset of phylogenetically related MDR isolates (SW070912, SW070910, SW071620, and PCP06238) as phylogroup D, suggesting that these isolates may be particularly concerning because of their potential to cause extraintestinal infections and to resist multiple classes of antimicrobials **(Figure 5, left, red text)**. Consistent with this notion, three of these isolates were identified by MLST analysis as belonging to the ST69 complex **(Figure 5)**, a pandemic *E. coli* lineage commonly associated with extraintestinal infections^36^, including urinary tract infections^37^. Interestingly, all phylogroup D isolates were from sewage (SW070912, SW070910, SW071620) or outfall (PCP06238), and all were isolated within the same three-week period (**Supplementary Table 3)**, thus directly implicating wastewater contamination in the emergence of MDR in the marine environment. Finally, all four isolates exhibited similar resistance profiles and plasmid-associated AMR gene content, via the disk diffusion assay and whole genome sequencing, respectively **(Figure 5)**, linking MDR phenotypes to specific AMR gene assemblages on plasmids^38^.

### Plasmid analysis of pathogenic *E. coli* demonstrates dynamic MDR

Given their pathogenic potential and large AMR burden, we turned our attention to the phylogroup D *E. coli* isolates SW070912, SW070910, SW071620, and PCP06238. The MDR plasmids in SW070912, SW070910, and PCP06238 are classified as conjugative, and the MDR plasmid in SW071620 is classified as mobilizable **(Supplementary Table 4)**. Conjugative plasmids are capable of self-transfer to other bacterial isolates, while mobilizable plasmids can transfer to other bacterial isolates but require the expression of other genetic factors to transfer^39^. We reasoned that the similar AMR phenotypes of these four *E. coli* strains may reflect congruent plasmids carrying the same AMR genes. To examine this more closely, we compared synteny between the four plasmids to understand the extent to which the AMR genes and the broader plasmid backbone are conserved between these isolates (**Figure 6A**). This analysis showed large regions of synteny between the three ST69 isolates, with the ST394 isolate (SW070912) showing more limited synteny **(Figure 6A, grey)**, and AMR genes often being contained within these syntenic regions.

**Figure 6:**
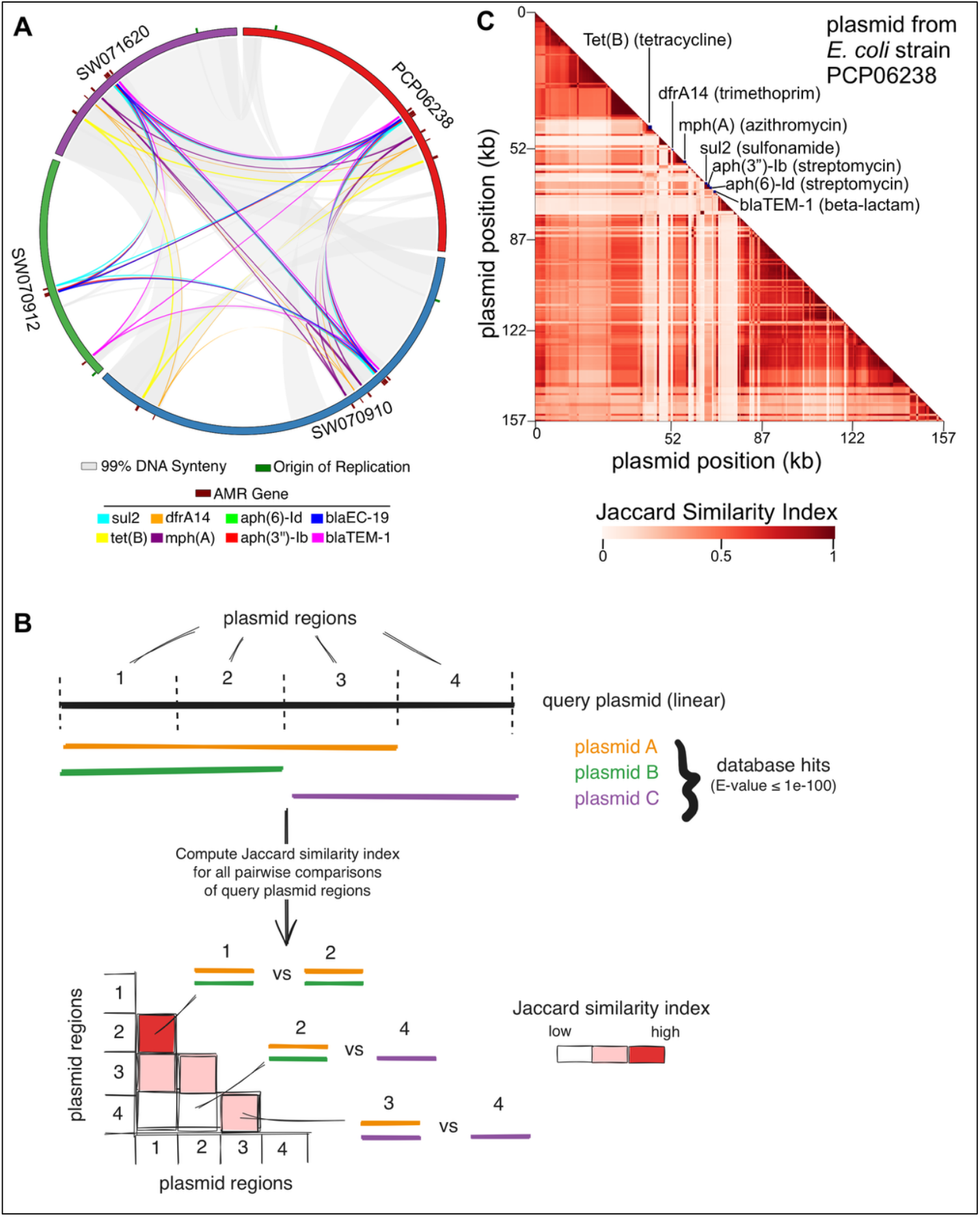
Plasmids from wastewater-derived Galápagos E. coli isolates harbor novel assemblages of AMR genes. (A) Circos plot showing plasmid and AMR gene synteny between 4 plasmids of interest. (B) Schematic illustrating how Galápagos *E. coli* plasmids were compared to plasmid database to identify and visualize conserved and novel regions as correlation heatmaps. (C) Correlation heatmap for the plasmid identified in *E. coli* strain PCP06238 (see Figure 5). Additional heatmaps are available in Supplementary Figure 3.

The high degree of synteny between the plasmids from ST69 isolates may simply be a consequence of the shared spatial and temporal origin of their parent strains. Conversely, since ST69 is a globally distributed lineage^38^ and the Galápagos is a site of intense international tourism, these plasmid sequences could be commonly observed in *E. coli* or other bacteria globally. Distinguishing between these interpretations is crucial because it could provide insight into whether potentially novel combinations of AMR genes are emerging in Galápagos because of wastewater contamination. To distinguish between these two interpretations of our data, we compared each of the four MDR plasmid sequences to the largest database of plasmid sequences created to date, spanning 40,757 globally distributed plasmids with significant counts of *Enterobacteriaceae*-origin plasmids^40^. Each plasmid was compared to the database via BLAST and database ‘hits’ were used to construct a correlation matrix depicted as a heatmap (**Figure 6B**, see methods). The color of each box in the heatmap represents the extent to which two regions of the same plasmid (X- and Y-axis) shared a common set of database hits (calculated as a Jaccard similarity) **(Figure 6B)**. This analysis and visualization identified novel recombinations of plasmid regions compared to existing plasmid sequences from the public database. For example, in plasmid SW071620 and SW070910, there is a ~20 kb segment with well-demarcated genomic borders **(Supplementary Figure 3A-B, respectively, arrows)**, indicating that this region is observed within the plasmid database (contiguous or non-contiguous). Nearly half of the ~100 kb plasmid SW070912 **(Supplementary Figure 3C, arrow)** shows high synteny to plasmids observed in the database, representing an example of a stable plasmid ‘backbone’. Annotating our correlation heatmaps with the location of AMR genes places resistance in the context of the conservation landscape for each plasmid. For example, plasmid PCP06238 **(Figure 6C)** – which originated from a phylogroup D MDR *E. coli* isolate that was recovered from ocean water and is closely related to wastewater isolates **(Figure 5)** – has seven AMR genes, all of which fall in regions of the plasmid that are neither commonly associated with the general backbone of the plasmid nor to other genetic regions carrying AMR genes. This phenomenon was also observed in the other plasmids examined from phylogroup D MDR strains isolated in the Galápagos **(Supplementary Figure 3)**, suggesting that AMR gene combinations leading to MDR were forming in plasmids without being linked to fully conserved cassettes or stable plasmid backbone lineages. Since bacteria carrying these plasmids were isolated from the environment or municipal wastewater with access to the marine environment, these plasmids have the potential lead to infections with pathogenic *E. coli* that are refractory for antimicrobial treatment and therefore have the potential to cause significant morbidity and mortality in animals or humans.

## Discussion

Our results implicate human wastewater contamination as a driving force for coliform contamination, environmental microbiome disruption, and the emergence of novel MDR *E. coli* in the marine ecosystem of the Galápagos, a UNESCO World Heritage Site cherished for its pristine natural beauty and rare animal species found nowhere else on the planet. Multiple sites – Carola pipe, Playa de Oro, the Muelle de Pescadores, and the Laguna (all either known or speculated sites of wastewater) – demonstrated strong human coliform contamination with differential temporal patterns of contamination (**Figure 2C**). These differences in coliform counts over time may be influenced by factors such as changing currents, weather patterns^41^, animal migration, or sewage release. Importantly, since none of our coliform assays are approved for guiding public health recommendations, it is impossible to know whether the levels of wastewater contamination we detected pose a significant threat to human health. However, tests conducted by local authorities in November 2024 (just four months after our last round of testing, shown in **Figure 2**), identified potentially dangerous levels of fecal coliforms and led the National Park of the Galápagos to temporarily restrict access to Playa del los Marinos and Playa de Oro^42^, the latter of which was also seen in our study to have relatively high levels of contamination on multiple occasions in 2023 and 2024, and as recently as July 2024 **(Figure 2C)**. Our results provide context for this policy decision and pinpoint specific locations, sources, and surveillance tools to monitor and intervene to minimize the threat of AMR bacteria to the biodiversity of this delicate ecosystem.

Genomic surveillance of AMR is critical to informing how and when resistance phenotypes may develop and move between human, animal, and environmental systems^43^. Our metagenomic sequencing **(Figure 3)** – guided by a ‘lab-free’ portable qPCR platform **(Figure 2)** – identified sites where the normal marine microbiome had been dramatically remodeled by wastewater contamination, resulting in the introduction of bacterial pathogens that included *Streptococcus, Bifidobacterium, Arcobacter, Pseudomonas, Clostridium, Bacteroides, Acinetobacter*, and *Escherichia*. The Galápagos is home to large populations of sea lions (*Zalophus wollebaeki*) and San Cristóbal in particular has some of the highest concentrations of sea lions in all of the archipelago^44^, with multiple rookeries (breeding grounds) and haulouts (resting areas) for this endangered species intermixed with tourist beaches, resulting in many close human-animal interactions^45^. Consequently, there is a high potential for wastewater-associated strains described in this study to colonize sea lions and lead not only to disease but also to further dispersal of AMR in the marine environment. *E. coli* is the most common bacterium associated with superficial wounds, skin abscesses, and ocular lesions in sea lions^46^ and has been linked to endocarditis^47^. Notably, higher rates of *E. coli* colonization and increased prevalence of AMR in sea lions and has been linked to proximity to humans^48,49^. The intense tourism and close intermixing of humans and sea lions on San Cristóbal raises the possibility that marine mammals could be at risk of exposure to MDR *E. coli*, underscoring the importance of viewing AMR as a One Health challenge, rather than simply a human health threat.

We observed significant AMR burden in our *E. coli* isolates, with a total of 64 MDR *E. coli* isolates that collectively exhibited 50 unique resistance phenotypes **(Figure 4)**, including three isolates that were resistant to meropenem, a carbapenem and ‘last resort’ antibiotic **(Figure 5)**. The World Health Organization recently designated carbapenem-resistant *E. coli* as critical priority pathogens^50^. The presence of so many unique MDR profiles in our data suggests that AMR within the Galápagos may be highly dynamic, with many gains and losses of AMR genes occurring through potential reassortment and recombination. Consistent with this notion, our comparative genomic analysis of plasmids from MDR *E. coli* showed that AMR genes appear to be subject to frequent ‘shuffling’ in plasmid backbones (**Figure 6** and **Supplementary Figure 3**). One interpretation of these data is that MDR in our *E. coli* isolates arises as a result of recent genomic changes within plasmids rather than simply through acquisition of extant MDR-conferring plasmids or other genomic regions. Therefore, plasmid reengagement/recombination offers one mechanism to explain much of the diversity and novelty of MDR phenotypes in the Galápagos (**Figure 4B**). The reassortment of genomic regions to generate novel MDR phenotypes in plasmids that may be highly mobile could lead to the introduction of MDR into pathogenic bacteria that ultimately lead to human or animal disease.

Although *E. coli* are most commonly associated with the lower gastrointestinal tract of mammals, they can also persist and even become naturalized into extraintestinal environments^51–55^, and coastal marine sediment has been previously demonstrated to be a reservoir for *E. coli* and a potential source for bacterial persistence and reemergence^56,57,58^. Once in the environment, AMR can spread rapidly. For example, within three months of the discovery of mcr-1, a plasmid-mediated colistin-resistance gene in Gram-negative organisms, it was detected in organisms in over 20 countries^59^. Within Latin America and the Caribbean, *E. coli* accounts for the highest pathogen-attributable fraction of human deaths associated with bacterial AMR^1^. This is of particular importance given our detection of multiple pandemic-associated *E. coli* strains, including ST10, ST58, ST69, and ST155 **(Figure 5)**. Curtailing the spread of AMR through ecosystems is vital to One Health but will require a deeper understanding of the mechanisms that lead to emergence of AMR strain and plasmids in complex microbial environments. Recent work with the healthcare-associated AMR plasmid pOXA-48 showed that horizontal transmission efficiencies can vary by orders of magnitude between different bacterial strains^60^. Using a population dynamics model, the authors went on to show that a single successful association of the plasmid with a bacterial host was sufficient to allow for establishment and stable maintenance in a complex microbial community and, ultimately, resistance to a broad range of antibiotic concentrations^60^. Similar experimental and modeling approaches with a focus on environmental – rather than gut-associated – microbiomes could help elucidate key factors that allow the dissemination of wastewater MDR plasmids though marine ecosystems.

Despite the clear threat that AMR poses globally to humans and animals, relatively little effort has been devoted to optimizing methods for AMR surveillance in LMICs where socio-economic factors can drive the rapid emergence of multidrug resistant organisms. Addressing this challenge will require creative solutions for dealing with limited access to modern laboratory infrastructure and unreliable electrical supply or cold chain. AMR research in the Galápagos poses additional unique challenges – with approximately 97% of the archipelago falling under the protection of national park (Parque Nacional Galápagos), removal of water, soil, and even DNA is highly restricted thus necessitating processing of samples on-site. We utilized a multi-pronged approach to investigate the impacts of wastewater on marine microbial diversity and environmental AMR burden in a major port town of the Galápagos islands. By focusing on field-deployable technologies and methods, we were able to carry out qPCR, culture-based AMR phenotyping, metagenomic sequencing, and whole genome sequencing, all without the need for access to a traditional laboratory. A major advantage to conducting all work on-site, rather than transporting samples to centralized laboratories, is that it affords the opportunity to engage regularly with community members and stakeholders. In addition, our use of portable, smartphone-powered qPCR technology and ready-to-use lyophilized qPCR reactions (Biomeme qPCR platform) and media pads for isolating bacteria eliminates time consuming and error prone procedural steps, lowering the barrier for engaging with high school- or college-level students without sacrificing the quality of research. Just as mobile surveillance efforts helped track Zika virus evolution in Brazil^61^, and decentralized sequencing proved vital for SARS-CoV-2 surveillance in Africa^62^, the approaches we outlined here provide a robust framework for community-driven AMR research in low-resource areas.

## Methods

### Sampling of wastewater, and marine and fresh water sources

Marine and freshwater samples were collected using sterile 210 mL Whirl-Pak bags (Whirl-Pak Filtration Group). Wastewater samples were collected using a modified Moore swab, as described previously^63^. Briefly, tampons were suspended individually from a string, lowered into the wastewater stream, and allowed to passively sample for one minute. The swab was recovered, sealed in a plastic bag, liquid was extruded from the swab by compressing the bag by hand, and the corner of the bag was cut to recover the liquid sample into a 50 ml conical centrifuge tube. Each sample’s collection time and geolocation were recorded using metadata from cell phone photographs taken at the time of collection. Samples were either processed immediately or stored at 4°C for ≤ 5 hours before processing.

### Water sample processing and portable multiplexed qPCR for fecal contamination

200 mL of each water sample was filtered through a 0.2 µm nitrocellulose filter in a 100 mL disposable analytical filter funnel (Nalgene) using a Mini LABOPORT N96 vacuum pump (KNF) or a hand-operated vacuum pump (Nalgene). Nitrocellulose membranes were recovered from filter funnels and placed in a Biomeme 5 mL sample homogenization tube containing a ½ inch steel ball bearing and 2 mL of lysis buffer. Material on the membranes was lysed by vigorous hand shaking for 1 minute. DNA was then extracted from 1 mL of homogenate using the M1 Sample Prep Cartridge Kit for High Concentration DNA (DNA-HI), per the manufacturer’s instructions (Biomeme). DNA was used immediately or stored at −20°C until use. To detect relative levels of human and animal fecal contamination, 20 µL of DNA was added to BioPoo Enterococcus Panel PCR Go-Strips (Biomeme), and qPCR was run on a Franklin ‘Three9’ thermocycler (Biomeme) controlled using the Biomeme Go app (v2.1.1) on a smartphone. The BioPoo assay is a triplex probe-based qPCR assay that simultaneously detects the following targets in each sample: 1) pan-stool marker, *Enterococcus faecalis* with probe conjugated to 6-carboxyfluorescein (FAM); 2) human-specific fecal marker, HF183 *Bacteroides* 16S rRNA with probe conjugated to Texas Red; and 3) Internal positive control (IPC) conjugated to cyanine5 (Cy5). Samples were considered positive if the CT value was less than 38. Freshly opened bottled drinking water was processed alongside ocean water and freshwater samples as a negative control in the qPCR.

### Metagenomic sequencing with Oxford Nanopore Technologies (ONT)

Material from ocean water samples was concentrated by filtering 2 L of ocean water through a 0.2 μm nitrocellulose filter in a disposable analytical filter funnel (Nalgene) using a hand-operated vacuum pump (Nalgene) or a Mini LABOPORT N96 vacuum pump (KNF). DNA was extracted from the filter using the DNeasy PowerSoil Pro Kit (Qiagen) with a modified preprocessing step. Briefly, the filter was placed in a Biomeme 5 mL sample homogenization tube containing a 0.5-inch steel ball bearing and 2 ml CD1 buffer. The tube was shaken by hand for one minute, and 800 ul of the homogenate was subsequently transferred to a PowerBead Pro tube. The remainder of the extraction proceeded per the manufacturer’s protocol. Sewage was extracted from 200 µl material using the DNeasy PowerSoil Pro Kit according to the manufacturer’s protocol. DNA as quantified using the Qubit dsDNA High Sensitivity Assay Kit (Thermo Scientific) and was used immediately or stored at −20°C. Sequencing libraries were prepared from 1 ug DNA using the Ligation Sequencing gDNA Kit (SQK-LSK110, Oxford Nanopore) with Solid Phase Reversible Immobilization (SPRI) beads. Libraries were sequenced with an Oxford Nanopore MinION using R9.4.1 flow cells.

### Analysis of ONT metagenomic data

Metagenomic long read data were basecalled using Dorado v0.3.1 using the super accuracy model (dna_r9.4.1_e8_sup@v3.6). The resulting fastq files were uploaded to the CZID online portal for long read metagenomic analysis^25^. Results were filtered based on nucleotide alignment length > 500, nucleotide percent identity > 85%, and nucleotide bases per million > 50. For AMR gene prediction, metagenomic data were processed as follows. Nanopore adapters were removed from reads with porechop (version 0.2.4), reads were quality filtered using filtlong (version 0.2.1), high-quality reads were assembled using Flye (version 2.9.5-b1801)^64^ with the meta flag, assemblies were polished with three rounds of racon (version 1.5.0) and two rounds of medaka (version 1.8.0), frameshifts were corrected using proovframe (version 0.9.7)^65^, and AMR genes were identified in contigs using AMRFinderPlus (version 4.0.19 with database version 2024-12-18.7)^26^. Heatmaps were generated in R.

### Fecal coliform and *E. coli* culture

1 mL of each sample was added to an MC Media Pad *E. coli* and Coliform test (Sigma) and incubated for 18-24 hours at 37°C. The next day, colonies of *E. coli* and other coliforms were counted by hand. Samples with growth too dense to be counted were diluted 1:10 (water) or 1:1000 (sewage) in sterile saline solution (Sigma) and re-cultured on a new MC Media Pad. Up to five *E. coli* colonies per MC Media Pad were subcultured on MacConkey II agar plates (Becton Dickinson) and incubated for 24 hours at 37°C for purity testing. Lactose-fermenting isolates were selected from MacConkey II plates with a sterile inoculation loop (Fisher Scientific) and suspended in 2 mL of sterile saline for antimicrobial susceptibility testing. The same inoculation loop was then submerged in 2 mL of LB broth, and the broth was incubated overnight at 35°C and 1000 rpm on a BioShake iQ thermomixer (QInstruments). Pellets from LB overnight cultures were stored at −20°C for DNA extraction and sequencing.

### Antimicrobial susceptibility testing

Kirby-Bauer disk diffusion assays were conducted to assess the antibiotic susceptibility of each isolate. Briefly, lactose-fermenting *Enterobacteriaceae* isolates or *E. coli* control stains ATCC 35218 and ATCC 25922 were visually adjusted to a 0.5 McFarland Standard (Hardy Diagnostics) in sterile saline solution. A sterile cotton-tipped applicator was then used to swab the bacterial suspension three times over the surface of two Mueller-Hinton II agar plates (Becton Dickinson) that had undergone manufacturer quality control analysis. After drying, six 6 mm antibiotic disks (Troy Biologicals) were pressed onto the surface of each plate. Twelve antibiotics were tested with each isolate: amoxicillin/clavulanic acid (20/10 mcg), azithromycin (15 mcg), cefoxitin (30 mcg), ceftriaxone (30 mcg), gentamicin (10 mcg), chloramphenicol (30 mcg), ampicillin (10 mcg), meropenem (10 mcg), streptomycin (10 mcg), tetracycline (30 mcg), sulfamethoxazole/trimethoprim (23.27/1.25 mcg), and ciprofloxacin (5 mcg). The zone diameters of clearance around each antibiotic disk were measured to the nearest millimeter after the plates were incubated at 37°C for 16-20 hours. When the full diameter could not be recorded (e.g., due to proximity to the plate edge), radii were recorded and doubled. For all antibiotics except for azithromycin, results were interpreted using zone diameter breakpoints (susceptible, intermediate, and resistant) according to the Clinical and Laboratory Standards Institute (CLSI) M100 standard^66^. Azithromycin was defined as wildtype or non-wildtype according to the CLSI M100 standard^66^.

### Whole genome sequencing and analysis for environmental *E. coli* isolates

Pelleted lactose-fermenting *Enterobacteriaceae* isolates were resuspended in residual LB broth, and DNA was extracted using the DNeasy PowerSoil Pro Kit (Qiagen) and quantified using the Qubit dsDNA High Sensitivity Assay Kit (Thermo Scientific). DNA was used immediately or stored at −20°C. Sequencing libraries were prepared from 400 ng DNA using the Native Barcoding Kit 24 V14 (SQK-NBD114.24, Oxford Nanopore). DNA was quantified after every step, and samples were normalized and pooled to equal concentrations for sequencing. Libraries were sequenced with an Oxford Nanopore MinION using R10.4.1 flow cells. Raw reads in POD5 format were basecalled using the Super Accuracy preset and demultiplexed using Dorado (version 0.9.6).

For analysis of isolate genomes, basecalled reads in fastq format were assembled with Autocycler (version 0.2.1)^67^ using flye (version 2.9.5-b1801)^64^, miniasm (version 0.3-r179)^68^, and raven (version 1.8.3)^69^ assemblers. Assembled contigs were re-oriented to the origin of replication with dnaapler (version 1.0.1). Anvio (version 8)^70^ was used to annotate each genome and construct a genome database for all isolates. Single copy genes (SCGs) were obtained using the anvi-get-sequences-for-hmm-hits function in Anvio using the Bacteria_71 set of SCGs. SCGs were then used to construct a phylogenetic tree with MrBayes (version 3.2.7)^71^ using concatenated protein output from Anvio. AMR genes were predicted with both NCBI AMRFinderPlus (version 4.0.19, with database version 2024-12-18.7 and the *Escherichia*-specific model)^26^ and Resistance Gene Identifier (RGI; version 6.0.4 with Comprehensive Antibiotic Resistance Database (CARD), version 3.2.7)^72^. Outputs from AMRFinderPlus and RGI were aggregated together by matching AMR gene names. MLST typing was done with mlst (version 2.19.0)^73,74^, Clermont phylogroups were determined using Clermontyping^75^, and plasmid mobility type was determined using MOB-suite (version 3.1.9)^76^.

We created a circos plot to visualize DNA and AMR gene synteny among four plasmids (PCP06238_contig_2, SW070910_contig_2, SW070912_contig_2, SW071620_contig_2). For plasmid synteny analysis, six pairwise alignments were performed using nucmer (MUMmer v4.0.1)^77^, regions were filtered to retain only significant matches (99% similarity), which were used to generated syntenic coordinates. For AMR gene synteny, coordinates of AMR genes were extracted from the results of AMRFinderPlus and RGI, and pairwise linkages were constructed for each AMR gene per plasmid. Plasmid and AMR gene synteny files were concatenated to create a links file for Circos (v0.69-8)^78^. Additionally, the origin of replication for each plasmid was predicted using OriV-Finder^79^ and annotated on the circos plot.

### Genomic Database analysis of *E. coli* plasmids

MDR plasmids from four *E. coli* isolates (SW070910, SW070912, SW071620, PCP06238; identified in sequencing as SW070910_contig00002, SW070912_contig00002, SW071620_contig00002, and PCP06238_contig00002, respectively) were analyzed against a database of 40,757 globally distributed plasmids^40^, integrated and deduplicated from the following databases: PLSDB (version 2020_11_19)^80^, Orlek^81^, COMPASS^82^, pATLAS^83^ and MOB-suite (version 3.0.1)^76^. Plasmids from the Galápagos *E. coli* isolates were compared to this database using command line BLAST 2.12.0+^84^ and all hits with an E-value ≤ 1e^-100^ were considered matches. For plotting BLAST results **(Figure 6C)**, each plasmid used to query the database was divided into 2000 equally sized regions. For each region, we collected the set of plasmid names (BLAST Subject IDs) from the global database BLAST hits to the region. This set of plasmid names was used as input for visualizing the pairwise Jaccard Similarity Index (J(A,B) = |A ∩ B|/|A ∪ B|) for different regions of the Galápagos plasmids. Identical sets of database plasmid BLAST hits to two regions within a Galápagos plasmid would yield a Jaccard index value of 1, and no common plasmids database plasmid BLAST hits to two regions would yield a Jaccard index value of 0 **(Figure 6B)**.

## Abbreviations

qPCR: Quantitative Polymerase Chain Reaction
AMR: Antimicrobial Resistance
MDR: Multi-drug Resistance

## Acknowledgements

We thank Fausto Rodriguez for facilitating access to the Galápagos National Park; Whitman Cox for providing guidance on wastewater management infrastructure and challenges on San Cristóbal island; and Demy Castillo for testing qPCR protocols on the island and providing feedback on assay usability. We also thank Jesse vanWestrienen and Max Pearlman from Biomeme, Inc. for guidance on qPCR assay selection and water collection and filtration strategies.

## Availability of data and materials

Metagenomic sequencing data and whole genome sequencing data for *E. coli* isolates are available on the Sequence Read Archive (SRA) at accession PRJNA1320681 and PRJNA1320685, respectively. All code used for analyses and figure generation is available on GitHub at https://github.com/ruicatxiao/GERA.

## Authors’ contributions

DPB and LMM designed and oversaw all experiments. AL, JCR, KV, MM, LE, MPJ, and KG carried out all field work and experiments, and participated in data analysis. AL and LMM generated and analyzed sequencing data. RX analyzed sequencing data and assisted with figure preparation. LMM, LE, and KK supervised undergraduate researchers during field work and assisted with all field work. SC assisted with the development, optimization, and interpretation of Kirby-Bauer disk assays. MW and EV managed programmatic aspects of the work and coordinated undergraduate recruitment, instruction, and mentoring. WC provided space and resources for temporary laboratory set-up. EV facilitated access to National Park sites and provided guidance on water collection sites and wastewater management on the island. DPB, LMM, and AL wrote the manuscript.

## Consent for publication

Not applicable.

## Supplementary tables

**Supplementary Table 1:** Listing of equipment and consumables that comprise the mobile AMR lab used for all work in this study.

**Supplementary Table 2:** Detailed information, including latitude/longitude coordinates and description, for all 20 sampling locations reported in this study.

**Supplementary Table 3:** Kirby-Bauer Disk Assay results for 183 isolates examined in this study. All measurements are in millimeters (mm) and resistance is coded as ‘r’ (resistant), ‘I’ (intermediate), or ‘s’ (susceptible) for all antibiotics except for azithromycin, which is coded as WT (wild type) or NWT (non-wild type). See methods section for more details on this assay and data interpretation.

**Supplementary Table 4:** Detailed information on all 155 *de novo* assembled contigs from *E. coli* isolate sequencing. The percentage of bases on each contig that were predicted to be modified by N6-methyladenosine (m6A), 5-methylcytosine (m5C), and N4-methylcytosine (m4C) is shown. GC content and coverage data from this table were used to generate plots shown in Supplementary Figure 2.

