## Supplementary figures and images for "Human wastewater contamination drives the emergence of novel multidrug resistant bacteria in the Galápagos marine ecosystem"

### Supplementary Figure 1

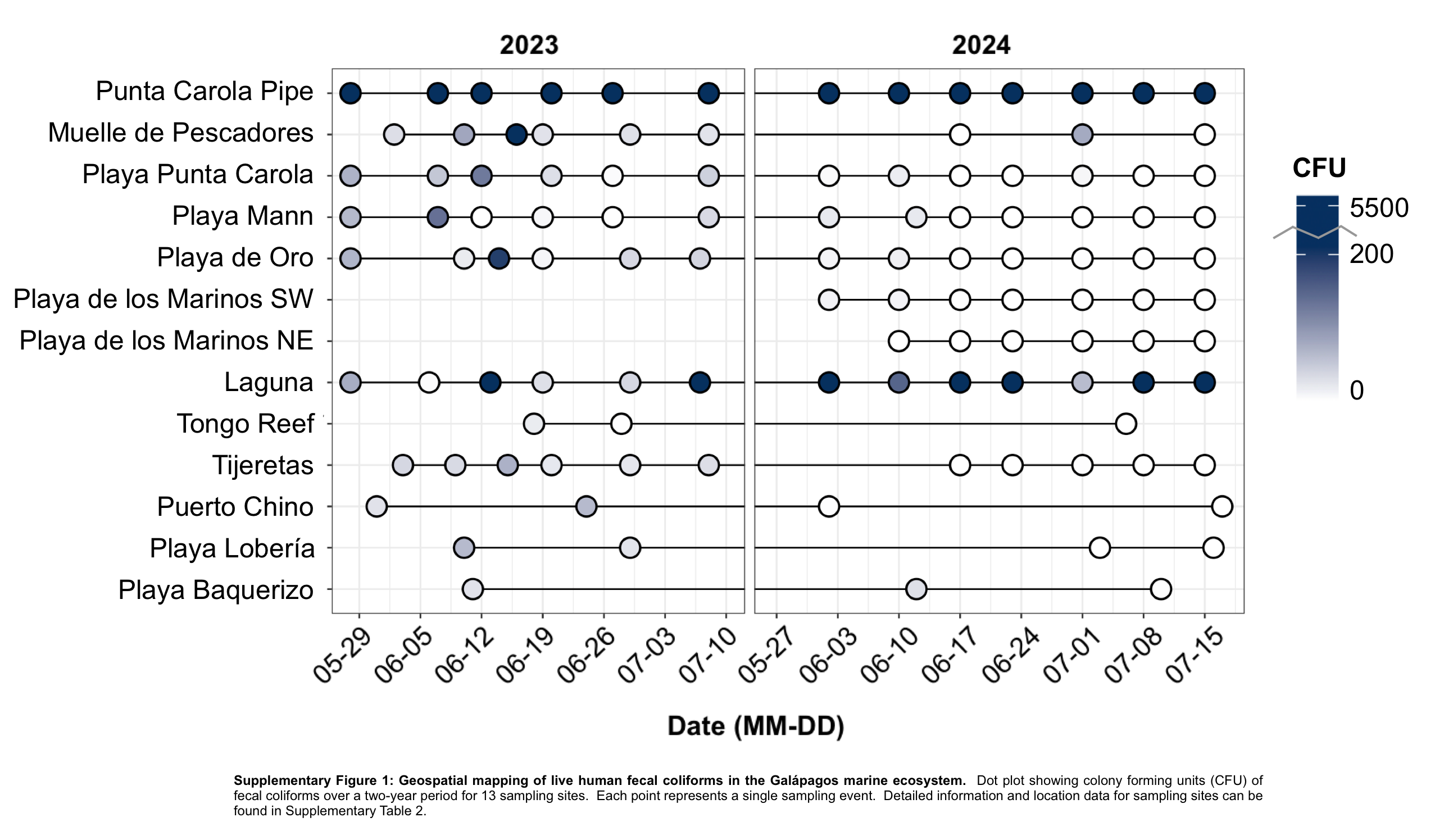

### Supplementary Figure 2

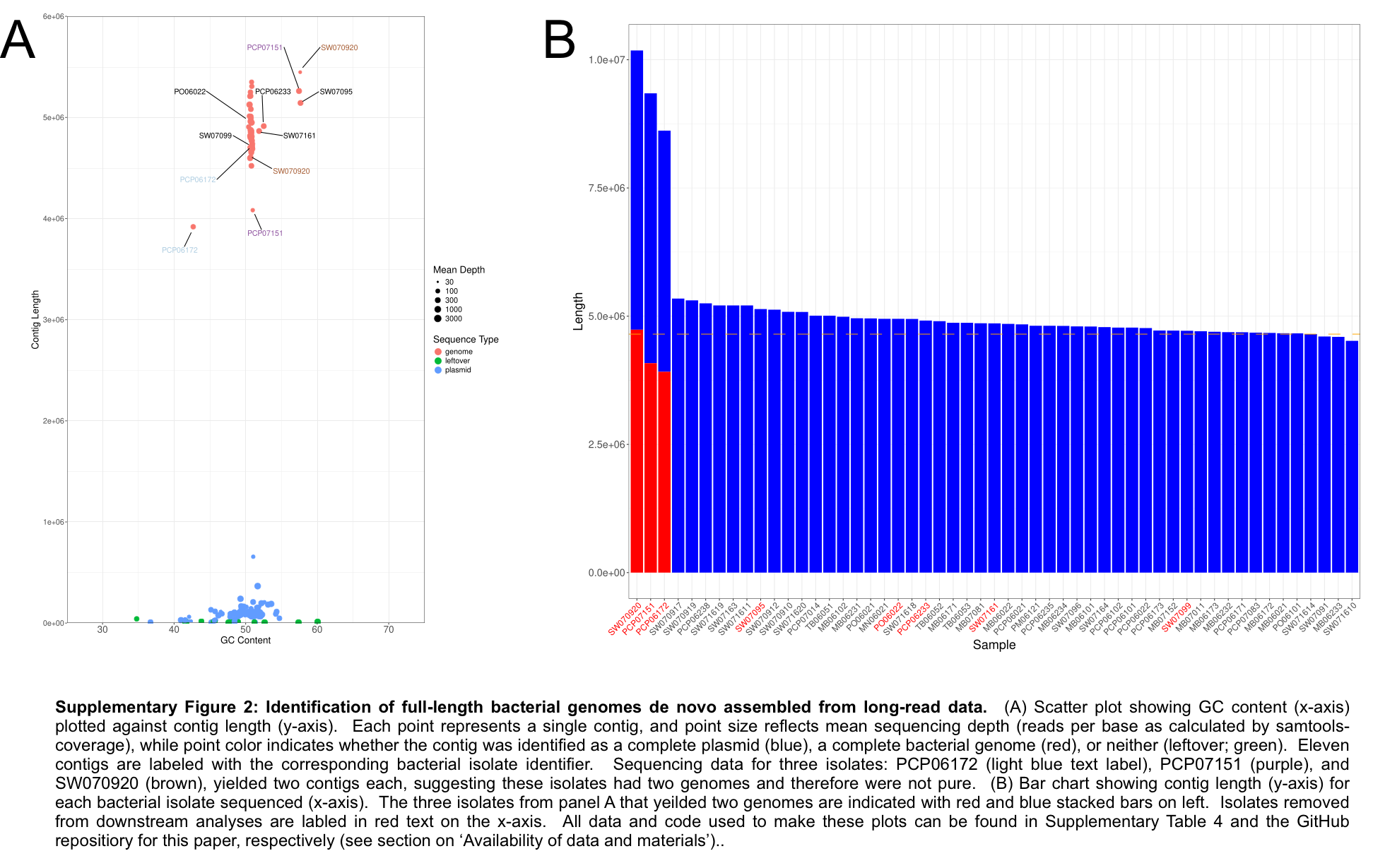

### Supplementary Figure 3

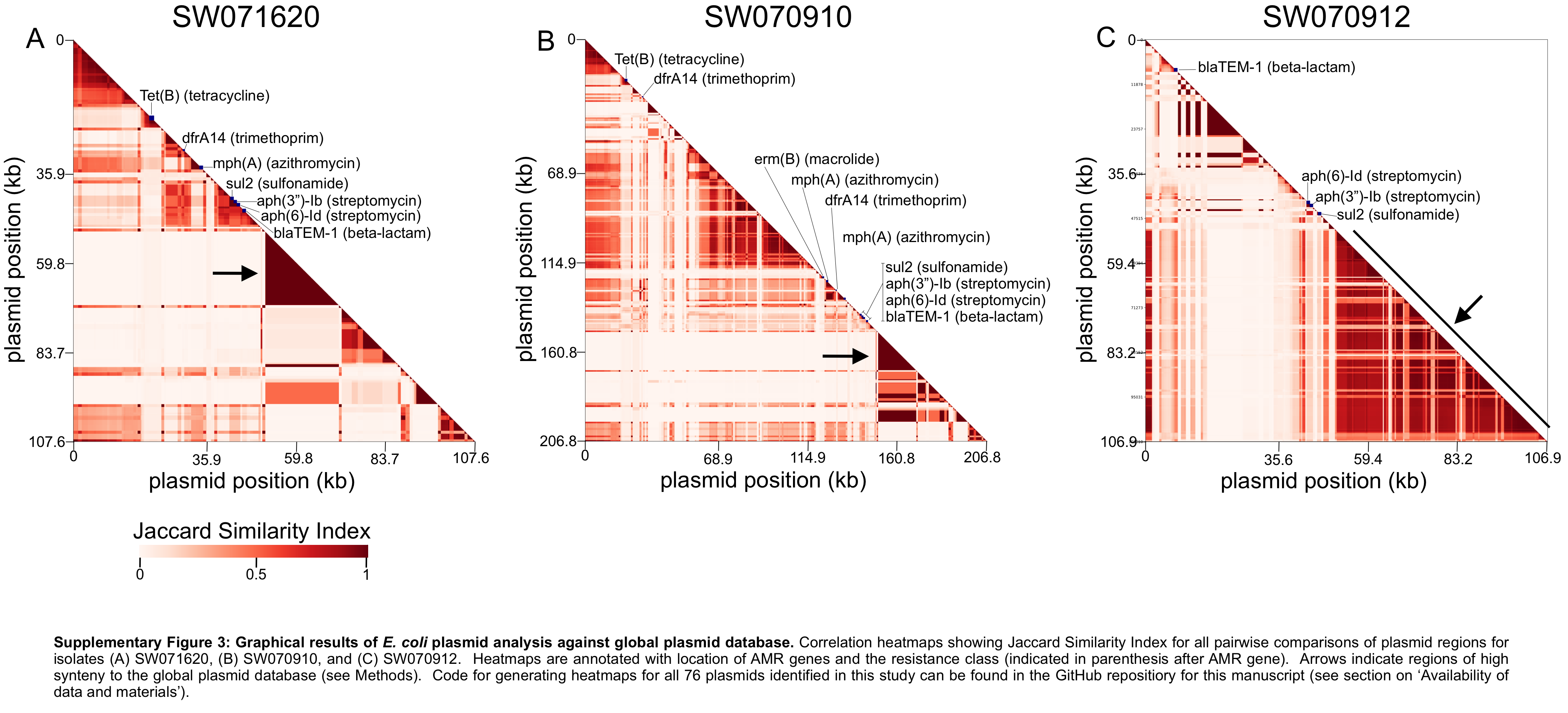
